# *In vitro* and *In vivo* Genetic Disease Modelling via NHEJ-precise deletions using CRISPR/Cas9

**DOI:** 10.1101/2020.03.25.007260

**Authors:** Sergio López-Manzaneda, Isabel Ojeda-Pérez, Nerea Zabaleta, Aída Garcia-Torralba, Omaira Alberquilla, Raúl Torres, Rebeca Sanchez-Dominguez, Laura Torella, Emmanuel Olivier, Joanne Mountford, Juan C. Ramírez, Juan A. Bueren, Gloria González-Aseguinolaza, Jose-Carlos Segovia

**Author notes:** Thérapie cellulaire et génique des maladies neurologiques de l’enfant et de l’adulte – NeuroGenCell, Equipe CARTIER, ICM - Institut du Cerveau et de la Moelle épinière, Hôpital Pitié-Salpêtrière, 47 boulevard de l’Hôpital, 75013 Paris, France Tfn. Corresponding author: Jose-Carlos Segovia, Differentiation and Cytometry Unit, CIEMAT (Building 70), Av. Complutense, 40, 28040 Madrid, Tfn.

## Abstract

Development of advanced gene and cell therapies for the treatment of genetic diseases requires confident animal and cellular models to test their efficacy and is crucial in the cases where no primary samples from patients are available. CRISPR/Cas9 technology, has become one of the most spread endonuclease tools for editing the genome at will. Moreover, it is known that the use of two guides tends to produce the precise deletion between the guides via NHEJ. Different distances between guides were tested (from 8 to 500 base pairs). We found that distances between the two cutting sites and orientation of Cas9 protein-DNA interaction are important for the efficiency within the optimal range (30-60 bp), we could obtain new genetically reproducible models for two rare disease, a Pyruvate Kinase Deficiency model, using human primary cells, and a (for *in vivo* primary hyperoxaluria therapeutic mouse model. We demonstrate that the use of a 2-guideRNAs at the optimal distance and orientation is a powerful strategy for disease modelling in those diseases where the availability of primary cells is limited.

## INTRODUCTION

Basic biology and the study of the function of the genome has been based on the availability of function deficient models, either cells or organisms, and the association of these loss-of-function with gene mutations. The availability of these models has allowed the research of human genetic diseases and, even, the development of gene therapy strategies for their treatment (Keeler, ElMallah, and Flotte 2017). The availability of endonucleases that can be directed to interact with very precise position in the cell genome has allowed the generation of knock-out models of any desired gene or position (Sakuma and Woltjen 2014; Silva et al. 2011; Deng et al. 2012).

Among all, CRISPR technology has become one of the most spread gene editing tools in the last years thanks to its easy design and its low cost. The action of these endonucleases produces the cleavage in both DNA strands in a precise manner according with the guide RNA (gRNA) position (Mojica and Montoliu 2016; Shibata et al. 2017; Nishimasu et al. 2014). The DNA cell machinery repairs these breaks either by non-homologous end joining (NHEJ), an error prone process, or by homology directed repair, that precisely corrects the damage. Both mechanisms have been extensively used to either eliminate the expression of a specific gene or to introduce new genetic material in a precise position of the cell genome, respectively. Repair by the non-homologous end joining machinery results in high variety of insertions, deletions (InDels) and sporadically inversions. This capacity to alter the original sequence has made these nucleases an excellent tool for the generation of knock out models, from bacteria to the human genome (Wassef et al. 2017; Islam 2018; Gilles, Schinko, and Averof 2015). Moreover, the deletion of specific regions that results in a recovery of function, either by eliminating a premature stop codon (Mettananda et al. 2017) or by deletion of specific gene regulators or silencers (Mojica and Montoliu 2016; Canver et al. 2014; Geisinger et al. 2016) has been suggested as an alternative therapy for specific diseases (Mettananda et al. 2017; Bonafont et al. 2019; Ousterout et al. 2015). In fact, a gene therapy clinical trial for the re-expression of the fetal globin in adult b-thalassemia patients by means of knocking out the BCL11A protein regulator is ongoing(“A Study to Assess the Safety, Tolerability, and Efficacy of ST-400 for Treatment of Transfusion-Dependent Beta-Thalassemia (TDT) - Full Text View - ClinicalTrials.Gov” n.d.).

NHEJ repairs double strand breaks in a non-predictive way, introducing InDels that can be very variable. Different reports have shown that the generation of two double strand breaks can facilitate the outcome of the NHEJ repair action(Canver et al. 2014; Zheng et al. 2017; Geisinger et al. 2016). Moreover, it has been proposed as a potential therapeutic option by eliminating mutated exons and recovering an almost normal although functional protein(Ousterout et al. 2015; Bonafont et al. 2019). Guo et al studied the efficacy of the NHEJ-precise deletion (NHEJ-PD) and how this process acts when guides are separated 23-148 apart, being the precise deletion of the DNA material between the two guide cuts the most common event(Guo et al. 2018). Thus, the use of two guides that could delete a defined DNA fragment and alter the open reading frame of the affected locus in an efficient and pseudo-controlled way could be used to generate KO models for the study of the biology of the cell or for the generation of cellular or animal models of rare diseases.

Here we reported the use of NHEJ-PD to generate cellular and animal models of genetic diseases. Moreover, we studied the optimal distances and the impact of the CRISPR-Cas9 machinery orientation in the NHEJ-PD.

## MATERIAL AND METHODS

### Cell lines

HEK293T cell line (human embryonic kidney cell line competent to replicate vectors carrying the SV40 T antigen; ATCC: CRL-3219) was cultured in Iscove’s modified Dulbecco’s medium (IMDM; Gibco), HyClone (10%; GE Healthcare) and penicillin/streptomycin (1%; Gibco). Cells were maintained at 5×10^5^ cell/mL concentration.

K562 cell line (chronic myelogenous leukemia; ATCC: CCL-243) was cultured in Iscove’s modified Dulbecco’s medium (IMDM; Gibco), HyClone (10%; GE Healthcare) and penicillin/streptomycin (1%; Gibco). Cells were maintained at 1×10^5^-1×10^6^ cell/mL.

Incubation conditions were the same for all cell lines used: 37ºC, 5% CO_2_ and 95% relative humidity.

### Animals

All experimental procedures were approved by the Ethics Committee of the University of Navarra and the Institute of Public Health of Navarra according to European Council Guidelines. All the experiments using animal models complied with all relevant ethical regulations. Agxt1-/- mice (B6.129SvAgxttm1Ull) were bred and maintained in a pathogen-free facility with free access to standard chow and water. Agxt1-/- mice were genotyped as described(Salido et al. 2006). Age-matched C57BL/6J mice (Harlan laboratories) were used as control animals. A final dose of 1×10^13^ vg/kg (5×10^12^ vg/kg/vector) was administered to 12-14-week-old Agxt1-/- male animals by intravenous injection. Animals were sacrificed and livers harvested 21 days after the administration of the vectors.

### Design and generation of CRISPR/Cas9 tools

For the experiments performed *in vitro* in cell lines, two different plasmids were used to clone specific guides. The plasmids used contained guide RNA and RNA scaffold expression driven under U6 promoter and also the Cas9-ZsGreen sequences linked by a P2A (porcine teschovirus-1’s motif that produce co-translational “cleavage” of both proteins) under the CMV promoter. The different guides were designed based on Zhang Lab’s program (MIT) against PKLR gene and chosen by score and its location in the gene.

The vectors and gRNAs used for the generation of a KO model in vivo were previously designed and individually characterized(Zabaleta et al. 2018)

### Human hematopoietic progenitor medium

Human hematopoietic progenitors were cultured in StemSpan SFEM I (Stem Cell Technologies) supplemented with human stem cell factor (hSCF, 100 ng/mL; EuroBioSciences), recombinant human thrombopoeitin (hTPO, 100 ng/mL; EuroBioSciences), human FMS-like tyrosine kinase 3 ligand (hFlt3-ligand, 100 ng/mL; EuroBioSciences), Glutamax (1%; Gibco), penicillin/streptomycin (1%; Gibco). Cells were maintained at 1×10^6^ cell/mL concentration. Incubation conditions were the following: 37ºC, 5% CO_2_ and 95% relative humidity.

### Erythroid differentiation protocol

Human hematopoietic progenitors were differentiated toward erythroid lineage using 4 different media. The first medium, days 1 to 7, was based on StemSpan SFEM I (Stem Cell Technologies) supplemented with hSCF (50 ng/mL; EuroBioSciences), hFlt3-ligand (16.7 ng/mL; EuroBioSciences), bone morphogenetic protein 4 (BMP-4, 6.7 ng/mL; Peprotech), human interleukin 3 (hIL-3, 6.7 ng/mL; EuroBioSciences), human interleukin 11 (hIL-11, 6.7 ng/mL; EuroBioSciences) and human erythropoietin (h-EPO, 1.3 U/mL; Amgen). Cells were then transferred to the second medium in which they were culture on days 7 to 14. Second medium was based on Iscove’s modified Dulbecco’s medium Glutamine supplemented (IMDM GlutaMAX supplemented; Gibco) enriched with BSA (1%; Sigma), insulin (0.01 mg/mL; Sigma), human transferrin (0.2 mg/mL; Sigma), β-mercaptoethanol (91 μM; Gibco), penicillin/streptomycin (1%; Gibco), lipid mixture 1 (1x; Sigma), ethanolamine (0.004%, Sigma), hSCF (5 ng/mL; EuroBioSciences), hIL-3 (6.7 ng/mL; EuroBioSciences), hIL-11 (6.7 ng/mL; EuroBioSciences), hEPO (1.3 U/mL; Amgen), insulin-like growth factor-1 (IGF-1, 20ng/mL; PeproTech) and hydrocortisone (1 μM; Sigma). After day 14 and for 2 days, cells were transferred to the third medium that was also based on IMDM GlutaMAX medium (Gibco) supplemented with BSA (1%; Sigma), insulin (0.01 mg/mL; Sigma), human transferrin (0.2 mg/mL; Sigma), β-mercaptoethanol (91 μM; Gibco), penicillin/streptomycin (1%; Gibco), lipid mixture 1 (1x; Sigma), ethanolamine (0.004%; Sigma) and hEPO (10 U/mL; Amgen). Final enucleation was carried out for five days on the forth medium that was also based on IMDM GlutaMAX medium (Gibco) supplemented with BSA (1%; Sigma), insulin (0.01 mg/mL; Sigma), human transferrin (0.2 mg/mL; Sigma), β-mercaptoethanol (91 μM; Gibco), penicillin/streptomycin (1%; Gibco), lipid mixture 1 (1x; Sigma) and ethanolamine (0.004%; Sigma).

All media were filtered by 0.22 μm before use. Sampling and analysis were done within this third week. During erythroid differentiation, cells were maintained between 4×10^5^ and 4×10^6^ cell/mL concentration. Incubation conditions were also 37ºC, 5% CO_2_ and 95% relative humidity.

### Erythroid differentiation immunophenotype

Immunophenotype of hematopoietic progenitors differentiated toward erythroid lineage was analyzed by flow cytometry using the following antibody combination: CD36-PE, clone CB38 (NL07), BD Pharmingen, 1μL ab/100μL; CD45-APCCy7, clone HI30, BioLegend, 1μL ab/100μL; CD71-PECy5, clone M-A712, BD Pharmingen, 1μL ab/100μL; CD235a-PECy7, clone HI264, BioLegend, 1μL ab/100μL). Cells were resuspended in a final volume of 100 μL, labeled for 30 minutes at 4ºC and washed with PBA (PBS with 0.2% sodium azide [Merck]; 1% BSA [Sigma]).DAPI (final concentration of 1 μg/ml) were added and events were recorded with a LSR Fortessa (BD Biosciences). Data were analyzed by FlowJo software (BD Biosciences).

### Electroporation of K562 cell line

DNA delivery was carried out on 20 μL volume format using Amaxa 4D-Nucleofector System and Cell line nucleofector solution SF (Lonza). Total amount of DNA electroporated was 3 μg (1.5 μg each plasmid in the case of using two plasmids at the same time). 2×10^5^ cells were electroporated following manufacturer’s protocol under FF-120 program.

### Electroporation of primary cells

Hematopoietic progenitor cells were electroporated with CRISPR/Cas9 plasmids as previously described, using Amaxa Nucleofector I (Lonza) and following manufacturer’s protocols. Cells were electroporated in cuvettes (electroporation volume of 100 μL) using Human CD34^+^ Cells kit (Lonza) and U08 electroporation program. Total amount of plasmid used was 11.26 μg, hence, when two plasmids were electroporated at the same time, 5.63 μg of each were added to the mix. After electroporation, cells were cultured on 24-well plate (FALCON) at 2 mL of final volume. In the case of the experiments with selection based on ZsGreen reporter expression, sorting was performed 24 hours after electroporation. Aggregates were eliminated before sorting using 40 μm Cell Strainer (FALCON).

### Deep sequencing analysis

Next generation sequencing (NGS) for a single cell suspension was carried out by STABVIDA and GeneWiz Companies. PCR products were treated and purified according to manufactures’s protocols. Samples were used for a library construction and sequenced with v3 chemistry in the lllumina MiSeq platform. Only variants represented in more than 1% were considered.

For deep sequencing of liver cells, total genomic DNA was extracted from the livers using NucleoSpin Tissue DNA extraction kit (Macherey Nagel) according to the manufacturer’s instructions. A first PCR round was performed using primers that amplified 390bp of mouse Hao1 gene (containing the targeted region in the middle of the amplicon) carrying the universal adaptor M13 sequence (underlined) in their 5’ end (forward 5’ <u>GTTGTAAAACGACGGCCAGT</u>AGACCAATGTTTGTCAGAGG 3’ and reverse 5’<u>CACAGGAAACAGCTATGACCT</u>AAAAGCATCCTAGGAAGGG 3’). Platinum Taq Polymerase (Invitrogen) was used. Subsequently, a second PCR was performed using primers targeting the universal M13 sequence and adding unique MID (multiplex identifier) barcodes to each sample. Sample pools were NGS-sequenced on the MiSeq Platfrom (Illumina, San Diego, United States) following manufacturer’s protocol.

## RESULTS

### The use of guide pairs allows precise deletion of the DNA sequences between the two induced double strand breaks

As a first attempt we studied if the use of two guides simultaneously could reducere the heterogeneity of the NHEJ activity to inactivate a certain gene. We choose the *PKLR* gene, responsible for the expression of the liver and the erythroid pyruvate kinase enzymes (LPK and RPK, respectively). Mutations in this gene are responsible of the rare disease Pyruvate Kinase Deficiency. Aiming to mimic mutations occurring in PKD patients, we selected exon 9 of the *PKLR* gene, the second exon with a highest number of mutations reported (33 different along the exon). Two guide RNAs were designed to cut in two hotspots of exon 9 where several mutations where grouped (Fig 1A; Fig EV1A).

**Figure 1.**
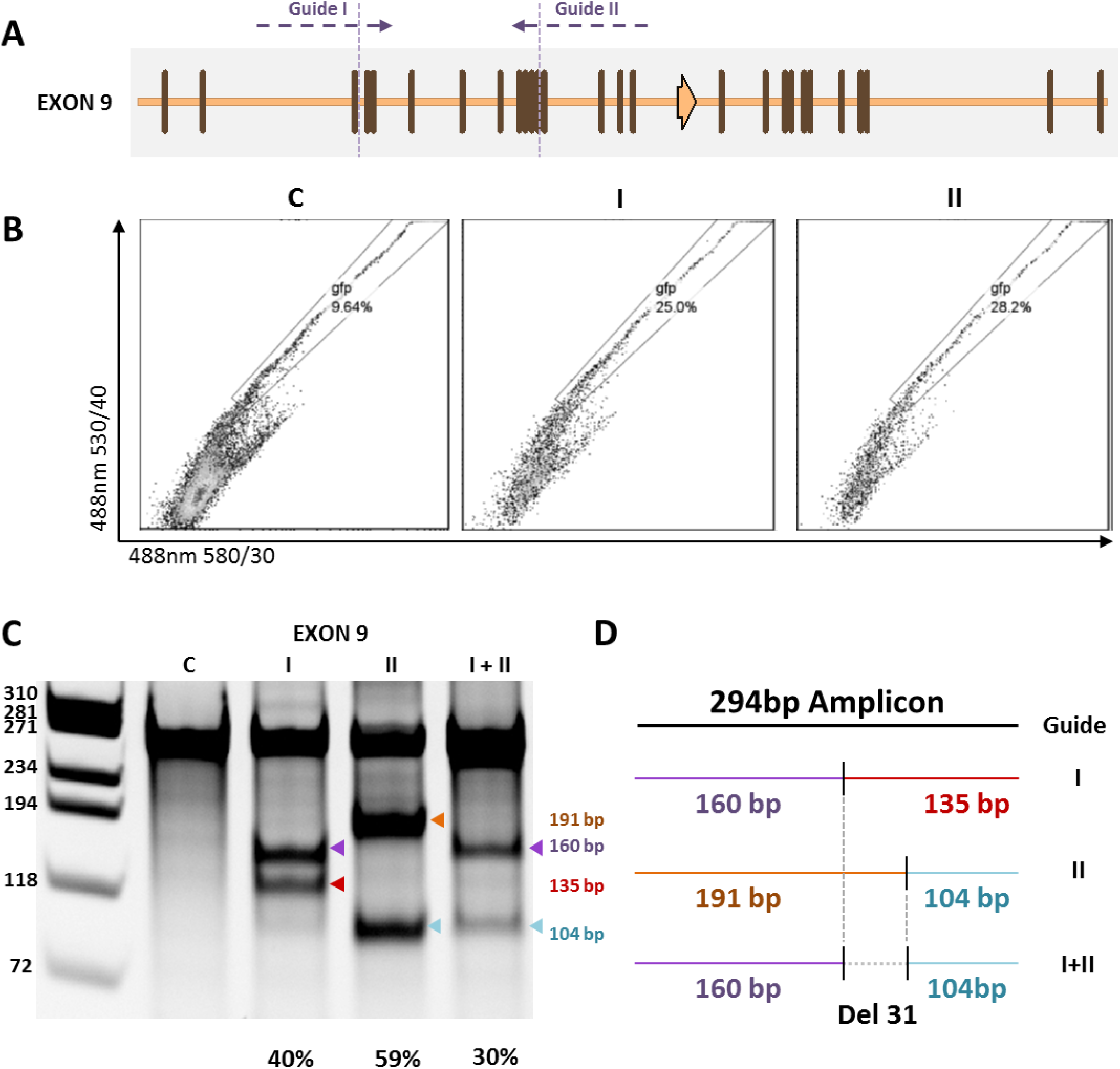
Design and use of CRISPR gRNAs in HEK293T cell line. **(A)** Exon 9 of the human *PKLR* gene is represented. Vertical lines and arrows are reported mutations, corresponding to one or more than one nucleotide affected, respectively. Horizontal dashed lines represent the two guide RNAs used. Vertical dashed lines represent predicted cut sites of guide RNAs selected. **(B)** Sorting of green fluorescence based on the ZsGreen present in the electroporated plasmids. 530/40 vs 580/30 nm dot-plot was used to distinguish ZsGreen fluorescence from auto fluorescence. The figure represent the ZsGreen fluorescence of the cell electroporated with the plasmid DNA containing only the Cas9 and the ZsGreen cDNAs (C), or containing the Cas9, the ZsGreen cDNAs and the sequences of the guide I(I) or the guide II (II). **(C)** Surveyor assay of the DNA obtained from the cells electroporated with the pasmids containing no guide, guide I, guide II or both guides together (I+II). The gene editing generate a characteristic band pattern for guide I (160 bp + 135 bp), for guide II (191 bp + 104 bp) and a mixed pattern when both are used at the same time (160 bp + 104 bp). **(D)** Scheme of the size bands that appear in (C) showing the 31bp deletion resulting after the simultaneous electroporation of guide I and II.

Human HEK293T cells were electroporated with all-in-one DNA plasmids containing sequences for one guide RNA, spCas9 nuclease and the green fluorescent protein (ZsGreen). Plasmids were electroporated individually or together. Seventy two hours after the electroporation cells were sorted based on ZsGreen expression (Fig 1B) and expanded up to confluence in 6-well plates. Surveyor assay was conducted in order to determine the activity of the designed guides and of the combination of both of them together. When guides were electroporated individually, the expected pattern of surveyor DNA fragments was obtained with high cutting efficiencies (40% for guide I and 59% for guide II; Fig 1C, 1D). When both guides were electroporated together, surveyor DNA fragments obtained were the combination of the cut of the two guides with the loss of the DNA fragment between the cut sites of the two guides (31 bp fragment). No evidence of fragments resulting from the cut of only one guide was observed (Fig 1C, 1D). These results confirmed previous evidences, demonstrating that the deletion of the DNA sequences between two guides, when used simultaneously, utilizing the CRISPR/Cas9 system, was highly frequent and the most common event of the NHEJ repair mechanism.

In order to define the joint between the two generated DSB we amplified the region of the joint and cloned it in PCR plasmids for their analysis by Sanger sequencing. All colonies analyzed that contained non-wt sequences presented the precise joint between the two DNA sides (NHEJ-PD; Fig EV1B).

### Distance between the two guides (30-500 bp apart) do not affect the generation of precise deletions

To test if the process was cell dependent, we performed a similar study using K562 (chronic myelogenous leukaemia cell line). Furthermore, in order to determine if the generation of the NHEJ-PD, being the most common event, was dependent of the distance between the guides used, five additional pairs were designed separated 30 bp to 500 bp, sharing always the upstream guide. Guide RNA pairs were designed to delete regions at exon 9, intron 9 and exon 10 of the *PKLR* gene (Fig 2A).

**Figure 2.**
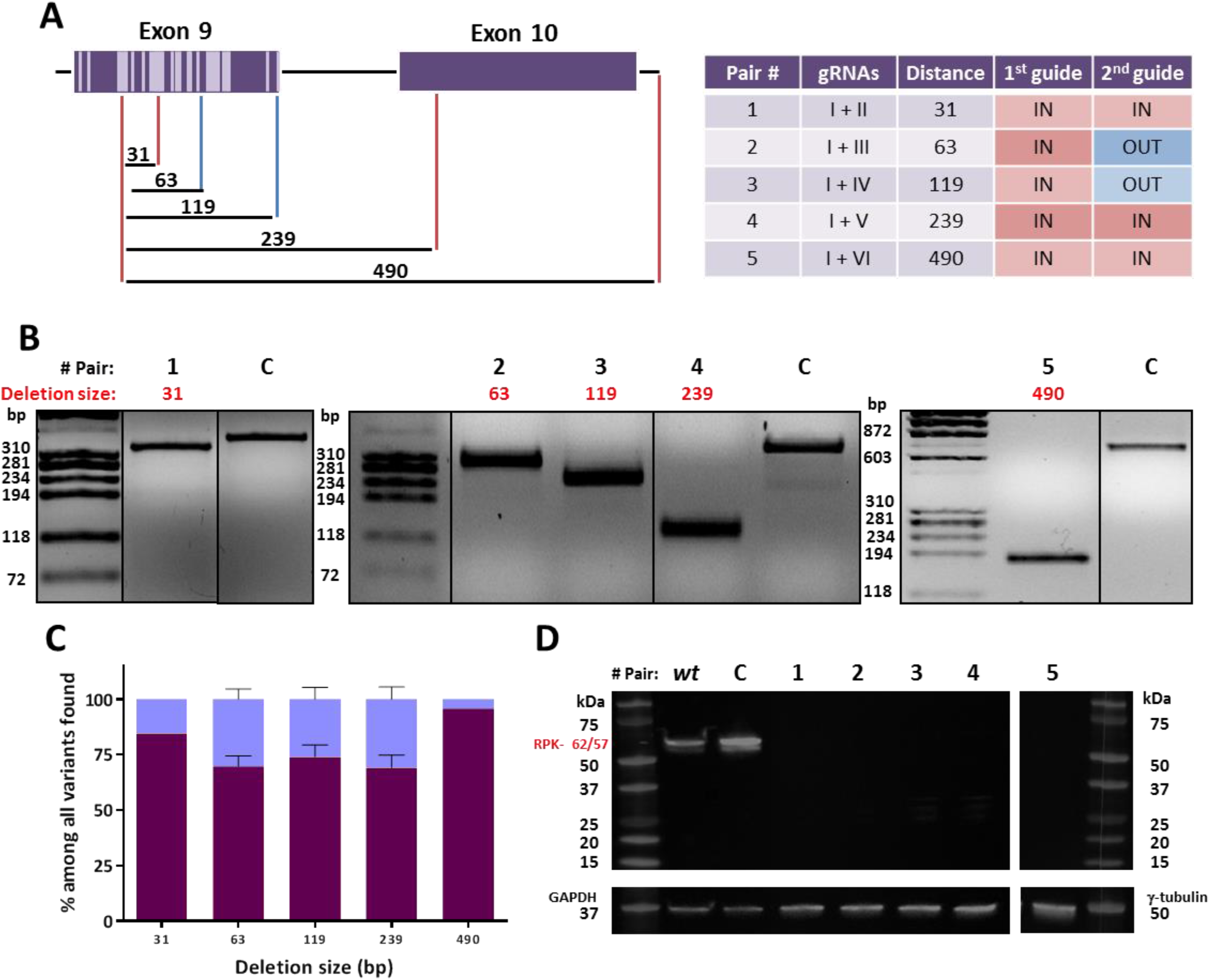
DNA and protein analysis of K562 cells electroporated with 5 guide pairs. **(A)** Left: Scheme of the 5 pairs tested. In dark purple: *PKLR* gene, exons are represented as boxes and introns as horizontal lines. Light purple represents mutations reported in exon 9. Vertical lines: position of the guides used, the different colors represent PAM position respect to the deletion, red lines: PAM “in” (inside the deletion sequence); blue lines: PAM “out” (outside the deletion sequence). Numbers above black horizontal lines are the size of the deletion generated in each combination. Right: list of the guides used, size of the deletion generated and its PAM disposition **(B)** PCR amplicons of the region of all combinations. Up: Pair number. Red: size of the precise deletion. C: control sample, K562 cells electroporated with Cas9-P2A-ZsGreen without guides and sorted. Pairs 1 - 4 were evaluated performing a 320 bp PCR meanwhile pair 5 was checked by a 700 bp PCR. **(C)** Next generation sequencing results from variants of the pairs tested, represented in more than 1%. Dark purple: precise deletion. Light blue: variants with modifications around expected deletion. Data are presented as mean ± SEM and Tukey’s Multiple Comparison statistical test was used to evaluate differences between groups. **(D)** Western blot of RPK pairs tested. Wt: non-electroporated K562 cells. C: control sample, K562 cells electroporated with Cas9-P2A-ZsGreen without guides and sorted. Black numbers: different pairs tested Red: sizes of RPK protein, complete (62 kDa) or after physiological N-terminal excision (57 kDa). GAPDH and γ-tubulin were used as loading control.

DNA plasmid constructs containing gRNAs and Cas9 were electroporated into K562 cells. Twenty-four hours post-transduction, cells were sorted based on the ZsGreen expression and expanded as described above. Editing efficacy was tested by PCR amplification of the affected region. Guide pairs showed a good capacity to delete gDNA, decreasing the size of the PCR product at the expected sizes. Indeed, wild type size products were not found (Fig 2B).

To deeply elucidate the outcomes that happened in the gene editing with the different guide pairs, PCR amplicons of the joint sequences were analyzed by next generation sequencing. Guide pair number 1, which precise deletion was predicted on 31 bp, presented only two events, being the most represented event the mentioned precise deletion with an 84.60% frequency. The other recorded event was also a deletion of 32 bp that was the same 31 bp deletion without an additional A at 5’ of the deleted sequence (precise deletion+1). Similar results were obtained with the other guide pair combinations (guide pair 2-63bp deletion: 69.70%; guide pair 3-119bp deletion: 74.05%; guide pair 4-239bp deletion: 69.18%; guide pair 5-490bp deletion: 95.84%) (Fig 2C). In all the other combinations, the remaining events, different from the precise deletion, were slight modifications surrounding it (Fig EV2A). When considering precise deletion plus precise deletion+1 together, the efficiency was always higher than 75% of the sequences analyzed in all the cases analyzed (Fig EV2C)

An additional guide pair with cutting sites separated 8 bp was also tested. Due to the proximity of the two cutting sites, the gRNAs were overlapping one each other. In this case, InDels detected where around one or the other cut site. No precise deletions or deletions containing the fragment between the cut sites were observed (Fig EV2B).

Finally, to corroborate the capacity of these deletions to knock-out the gene, a western blot assay of the edited populations was performed. No RPK protein was detected in any of the edited samples, indicating that any combination of guides was capable of eliminating completely the production of the protein (Fig 2D).

### PAM orientation of the guides affects the efficacy of precise deletions

One of the potential reasons to explain why a short distance between the two guides is less efficient in generating precise deletions could be the difficulties to access the two cutting sites at the same time due to allosteric or space impediments. To study these potential difficulties, we designed and cloned gRNAs spaced approximately 30 bp that facilitated the Cas9-DNA interactions in all 4 possible orientations, PAM-in/PAM-in (or-1), PAM-in/PAM-out (or-2), PAM-out/PAM-in (or-3) and PAM-out/PAM-out (or-4) (Fig 3A). Human K562 cells were electroporated with pairs of all-in-one DNA plasmids containing sequences for one gRNA, spCas9 nuclease and ZsGreen. Twenty four hours after electroporation cells were sorted based on ZsGreen expression and expanded as above. Total DNA was extracted from the 4 different combinations. Three different experiments were conducted with each combination. NGS was performed and bioinformatics analysis was performed to define the incidence NHEJ-PD and additional InDels in the genomic region. The combination or-1 (PAM-in/PAM-in) showed the best NHEJ-PD efficiency and the or-4 combination (PAM-out/PAM-out) was the worst one, including a high proportion of non-edited sequences. Combinations or-2 (PAM-in/PAM-out) and or-3 (PAM-out/PAM-in) were also efficient but the percentage of InDels different from the precise deletion were higher (Fig 3B and Fig EV3A). Interestingly, if we analyze the cases where only one guide cut, the guide oriented PAM-in was always more efficient (data not shown). As observed above, when we considered together the NHEJ-PD plus the NHEJ-PD+1 the percentage of edited sequences were close to 100% in the or-1 combination (PAM-in/PAM-in) (Fig EV3B). Altogether, these experiments showed that steric difficulties could be taking place in the efficacy of the cutting activity of the guides, suggesting that orientation of the PAM and positioning of the Cas9 nuclease on the DNA are important in the generation of short deletions by NHEJ.

**Figure 3.**
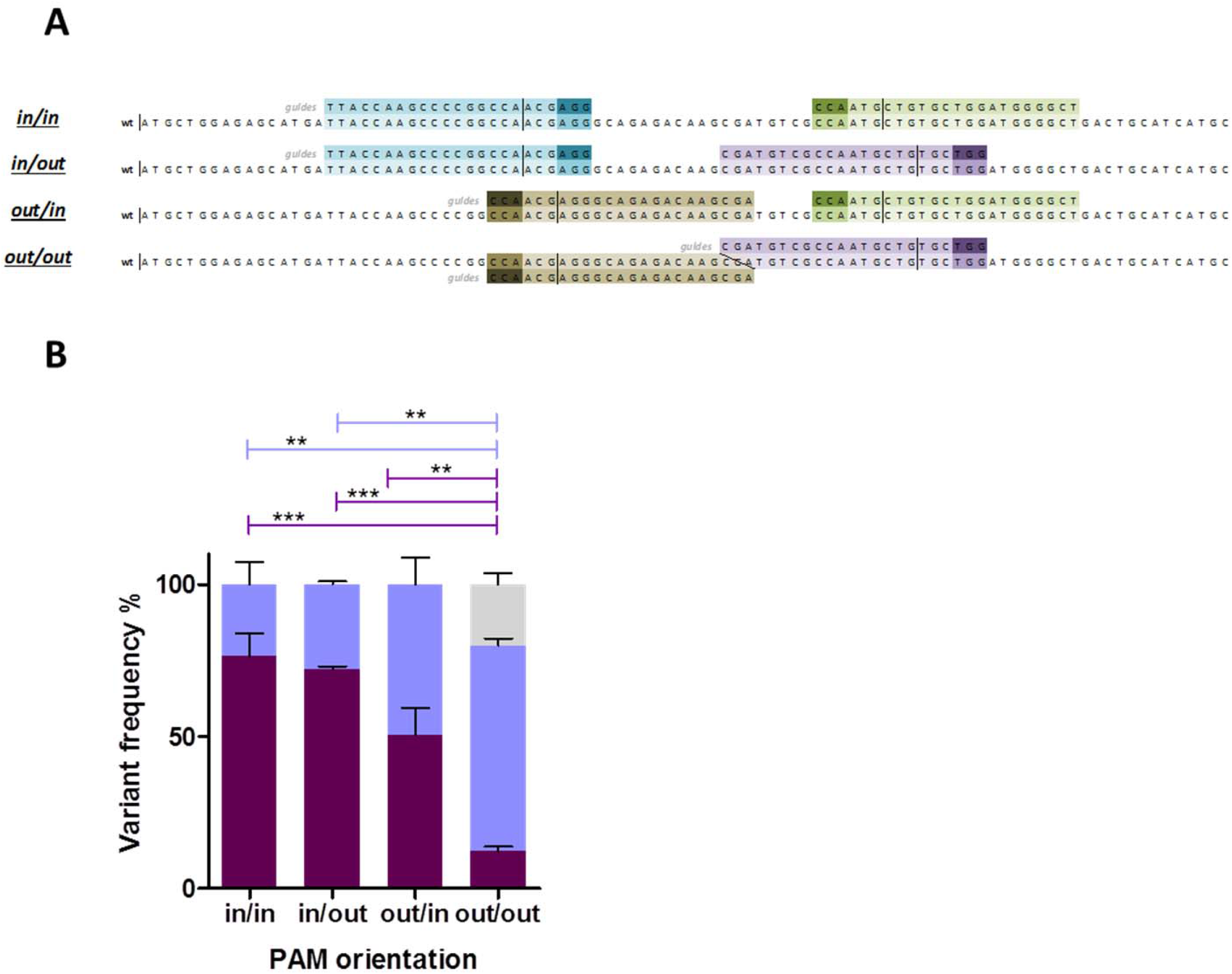
DNA analysis of K562 cells electroporated with 4 guide pairs separated 28-34pb apart providing all the PAM-In and PAM-out combinations. **(A)** Scheme of the 4 different combinations tested. In light colors: guides. In dark colors: PAMs sequences. The orientations of the PAMs are represented on the left, being “in” when the PAM is inside of the precise deletion and “out” when it is outside. **(B)** Next generation sequencing results from variants of the different combinations, represented in more than 1%. Dark purple: precise deletion. Light purple: variants with modifications around expected deletion. Grey: wild type. Data are presented as mean ± SEM and Tukey’s Multiple Comparison statistical test was used to evaluate differences between groups.

### NHEJ-PD generates a cellular model of PKD-like CD34 primary cells

Next, we wanted to test if the NHEJ-PD would also occur in primary cells and if this mechanism could be used to generate cellular models of genetic diseases in primary cells. For that, we applied the guide pair 1 to knock-out the PKLR gene in human cord blood CD34+ (CB-CD34+) cells. CD34+ cells are responsible for the generation of all the hematopoietic lineages, including the erythroid lineage where RPK expression is required for the proper production of mature erythroid cells. Reduction or lack of RPK function produces Pyruvate Kinase Deficiency (PKD). Healthy CB-CD34+ where electroporated, sorted and differentiated towards erythroid lineage. Afterwards, the modified region was analyzed by deep sequencing. Similarly to the results obtained in immortalized cell lines, NHEJ-PD was consistently achieved in edited CB-CD34+ cells, easily traceable following the decrease of the size of the PCR product (Fig 4A). Furthermore, the proportion of precise deletions was followed by the analysis of the cleavage produced by BstXI which target sequence is generated after precise deletion and is not present in the wild type neither in other deletions such as the 32 bp one (Fig 4B). Interestingly, the most represented events recorded in this primary cells were the same found in the cell line studies (31 bp deletion: 82±2 %; deletion 32 bp: 16±4 %). In addition, in one of the samples two other variants were found, representing around 1.5% of the events. Both were deletions, one of 40 and another of 55 bp, also centered in the precise deletion region (Fig 4C). Thus, the use of this two selected guides demonstrated the capacity of consistently limit the editing events to just this two that, afterwards, generate stop codons (Fig 4D).

**Figure 4.**
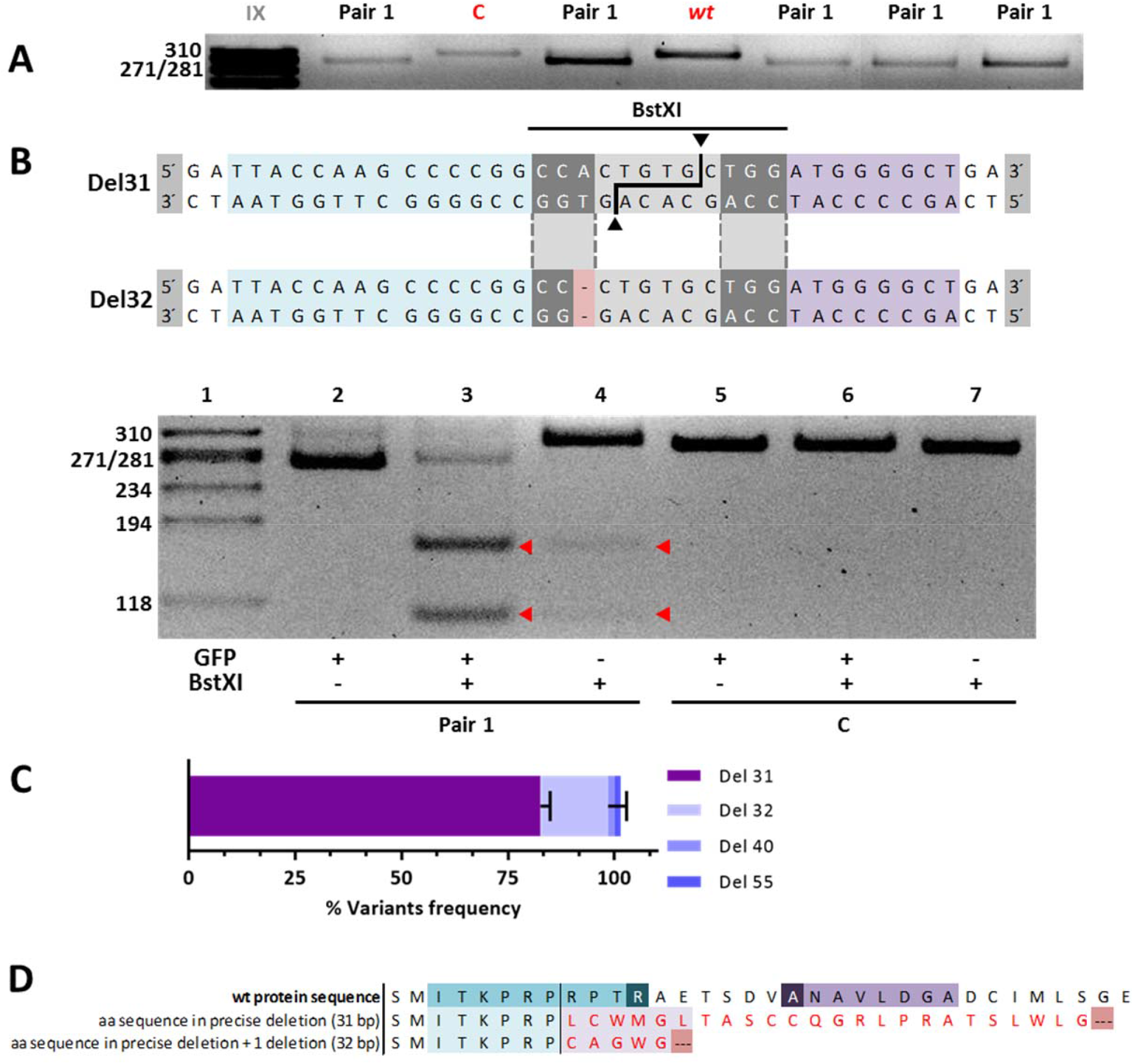
Primary CD34+ cells modified by Pair 1 gRNAs and differentiated towards erythroid lineage. **(A)** Exon 9 PCR amplicons from different experiments of hematopoietic progenitor cells differentiated *in vitro* toward erythroid lineage and edited with guides I + II (Pair 1). C: control cells electroporated with Cas9-P2A-ZsGreen without guides. I + II and C are the ZsGreen^+^ fraction selected by sorting. Wt: non-electroporated cells. Pair 1: edited cells from different experiments. **(B)** Up: Sequences resultant from the two main deletions produced after gene edit. Blue: region covered by guide I. Purple: region covered by guide II. Sequence in grey: BstXI recognition site. Light red: differential nucleotide, not present in 32 bp deletion, that is required for BstXI recognition. Down: Lane 1, molecular weight reference. 2-4 lanes: hematopoietic progenitor cells differentiated *in vitro* toward erythroid lineage and edited with guides I + II. 5-7 lanes: control cells electroporated with Cas9-P2A-ZsGreen without guides. ZsGreen, + or – fractions separated by sorting. + or – BstXI, treated or not with the enzyme. Red triangles, DNA cut after enzyme digestion. **(C)** Next generation sequencing results from two edited DNA samples, colors represent each of the deletions found named Del 31, Del 32, Del 40 and Del 55. **(D)** Amino acid sequence resultant after 31 and 32 bp deletions. Red amino acids are the alerted ones with respect to the wt. Both deletions generate premature stop codons represented by dots on red background.

In order to check if the edited cells reproduced the *in vitro* PKD phenotype, CD34-PKLR^-/-^ cells were cultured in specific media to allow erythroid differentiation. After a two-week differentiation protocol, cells were analyzed in terms of cell morphology, immunophenotype and pyruvate kinase activity. Non-edited CB-CD34+ cells and bone marrow CD34 cells obtained from a PKD patient were also differentiated following the same protocol. No differences were observed in the *in vitro* differentiation of the different samples demonstrating that PKD progenitors differentiate normally *in vitro* under the conditions used (Fig 5A). Similarly, analysis of cytospins performed at the end of the culture corroborated also the same degree of differentiation in all of the groups, independently of its deficiency in PKD, having the orthochromatic erythroblast population as the most represented cell type (Fig 5B).

**Figure 5.**
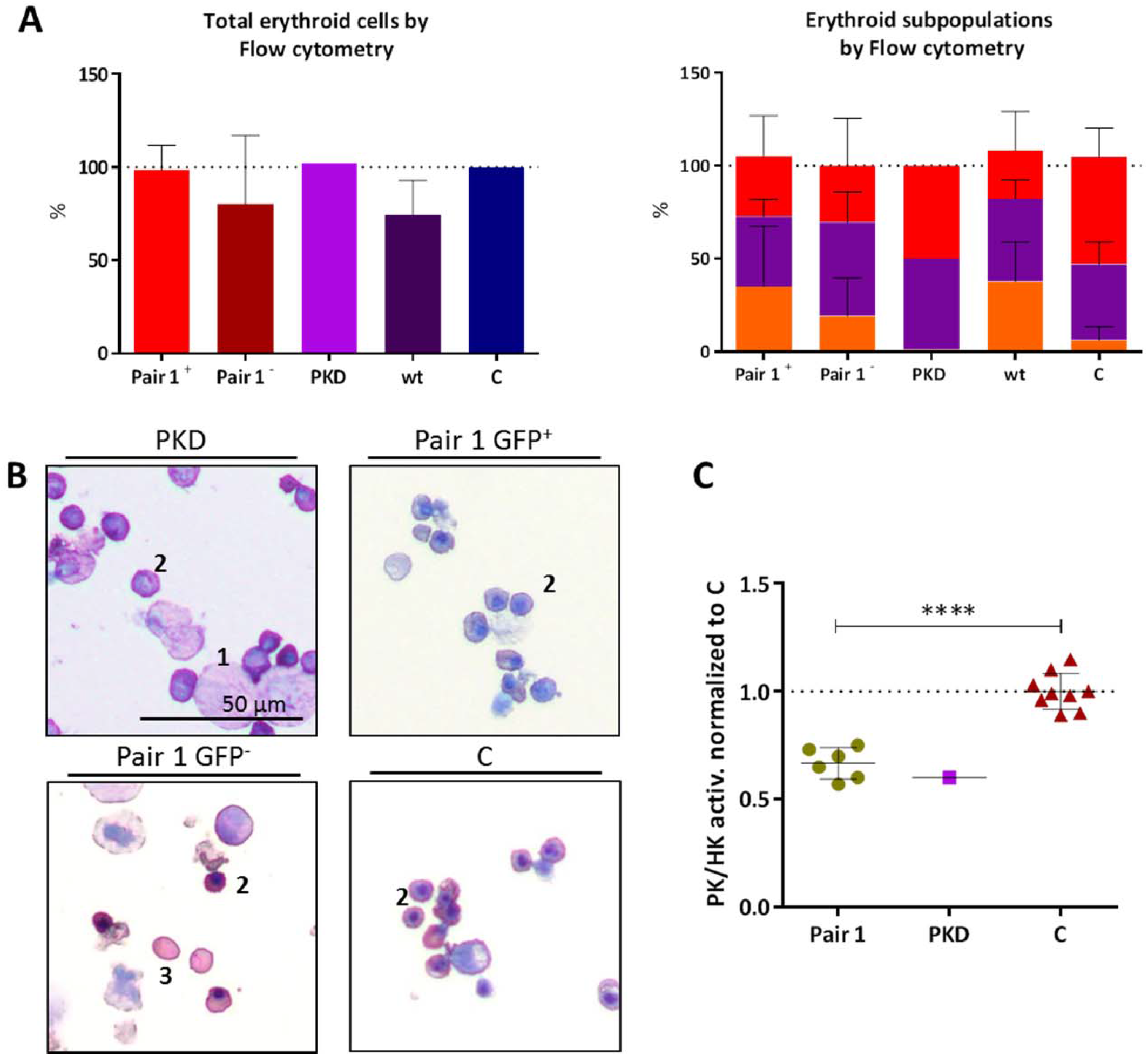
Immunophenotype and activity of hematopoietic cells edited and differentiated towards erythroid lineage. **(A)** Left: Percentage of cells considered erythroid based on their CD36/CD45 staining (it includes cells in transition from CD45^+^/CD36^+^ to CD45^-^/CD45^-^; values normalized with respect to control samples). Right: erythroid maturation followed by CD71/CD235a staining. Immature: CD71^+^/CD235a^+^ (orange); mature: CD71^-^/CD235a^+^ (red); transition from immature to mature: (purple). (n=6). PKD: cells differentiated from patient bone marrow sample. Pair 1: cells electroporated with guides I + II. C: control cells electroporated with Cas9-P2A-ZsGreen without guides. + or - : ZsGreen positive or negative fraction separated by sorting. Wt: non-electroporated/sorted cells. **(B)** Cytospin after 14 days of erythroid differentiation. 1: polychromatophilic erythroblast; 2: orthochromatic erythroblast; 3: reticulocyte. **(C)** PK/HK activity of edited cells after erythroid differentiation. Green circles: ZsGreen^+^ fraction cells electroporated with the two guides from different experiments. Purple square: PKD cells differentiated from patient bone marrow sample. Red triangles: control cells. Measures are normalized with respect of PK/HK of control samples, which were consider =1 (**** P<0.0001; n=6)

Finally, edited, control and PKD patient cells were tested for pyruvate kinase activity. PKD activity was tested together with the hexokinase (HK) activity, to normalize the increased PK activity of the most immature erythroblasts. Ratio PK/HK of the edited cells showed a decreased enzymatic activity comparable with the values obtained in the PKD patient sample differentiated in parallel and, approximately, 0.6 times the values of the healthy control cells (Fig 5C). Overall, NHEJ-PD after Cas9 DNA breaks by means of the use of guide pair combination generates an *in vitro* cellular model, resembling the characteristics of patient deficient cells, which would be very valuable for further testing alternative therapeutic strategies.

### NHEJ-PD generates *in vivo* knock-out animal model

In order to study if the use of two closed guides to minimize the variability of NHEJ could also work *in vivo*, we addressed the knock-out the mouse *Hao1* gene, which encodes for the glycolate oxidase (GO), in the liver of mice injected with AAV vectors expressing 2 different gRNAs and the *Staphiloccocus Aureus* Cas9 protein. Both gRNAs targeted opposite strands of exon 2 of mouse *Hao1* gene, were oriented in PAM-out orientation and DSBs were 64bp apart (Fig 6A). NGS analysis of the livers of animals treated with both vectors revealed that inside the edited cells, the majority of the sequences presented the deletion of the 64bp harbored between the two DSBs (Fig 6B). Moreover, these genetic modifications led to a decrease in *Hao1* mRNA levels (Fig 6C) and to the complete elimination of GO protein analyzed by WB (Fig 6D).

**Figure 6.**
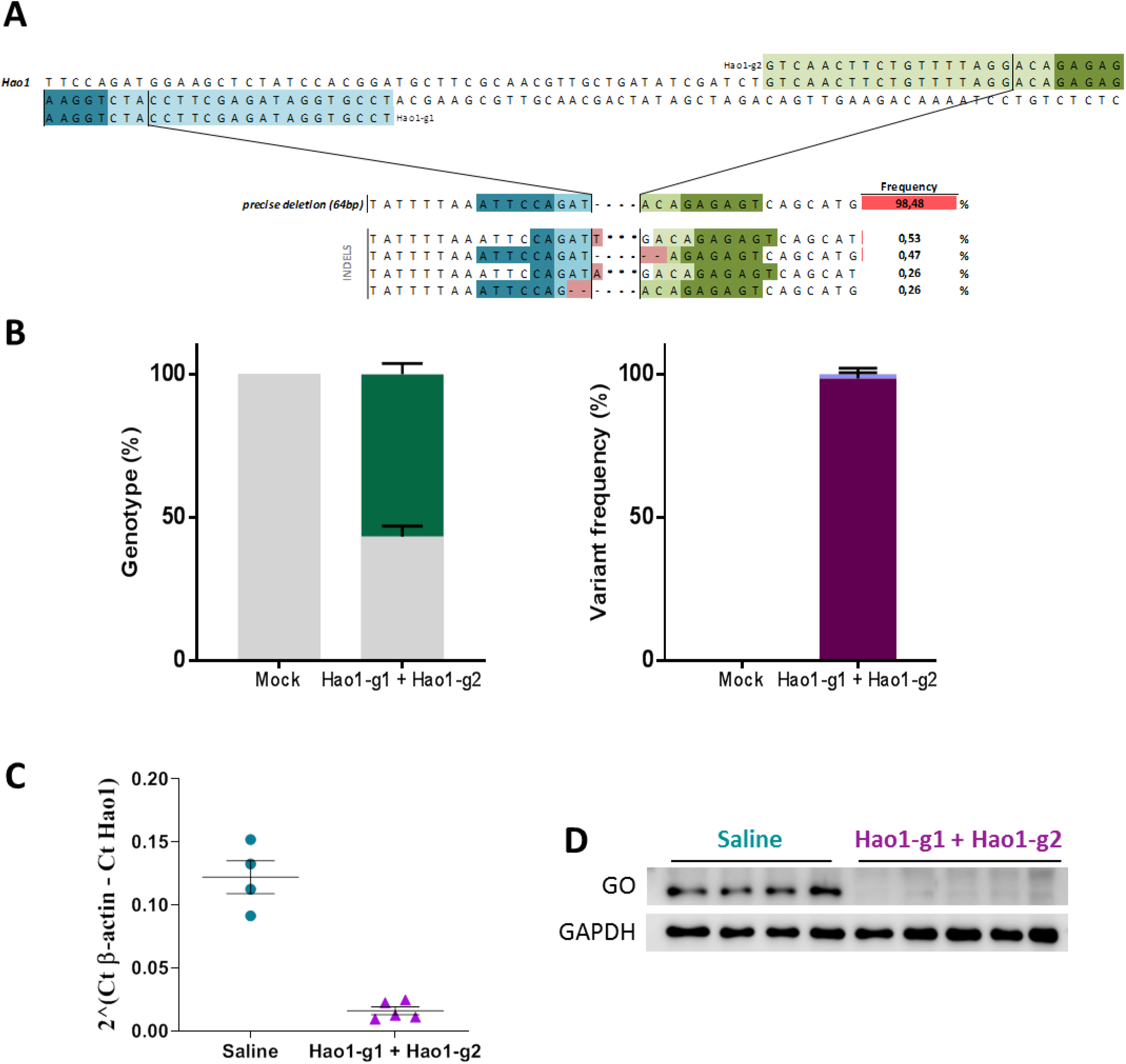
*In vivo* generation of NHEJ-PD generates a Hao1 gene knock-out in mice. **(A)** Scheme of the two guides tested in Hao1 gene. In light colors: guides (Hao1-g1 and Hao1-g2). In dark colors: PAMs sequences. Within the pair scheme there are three differentiated parts. Up: wt sequence and the guides used. Middle: precise deletion sequence and its frequency. Bottom: InDels near precise deletion and its frequency. NGS was performed for two different groups of mice, ones treated with a saline solution, and the others treated with the two guides and the SaCas9. Stars represent the wild type sequence and scripts are deletions respective to the wild type sequence (**B)** Next generation sequencing results from variants of the different combinations. Left: genotype of the different groups. In grey: WT mice; in green: edited mice. Right: percentage of the different edition variant; Dark purple: precise deletion; Light purple: variants with modifications around expected deletion. **(C)** Quantification of Hao1 mRNA expression levels by RT-qPCR in animals treated with saline and the combination of Hao1-g1 and Hao1-g2. **(D)** Western blot analysis of GO protein levels in representative animals treated with saline and the combination of Hao1-g1 and Hao1-g2. GAPDH was used as loading control.

## DISCUSSION

Herein, we have shown that the use of two guide RNAs to drive Cas9 cutting in adult cells facilitates the precise deletion of the sequences between them and reduces, or even eliminates, the heterogeneity of the error-prone NHEJ, allowing the knock down or knock out of the desired gene and the generation of cellular or animal models, with an almost homogeneous genotype. Previous works have indicated the preference for the NHEJ repair by the precise deletion between the two DSB generated when two guides are used simultaneously(Guo et al. 2018). Here, we demonstrate that this characteristic facilitates the generation of cellular and animal models with a more homogeneous and controlled genotype modification.

We have observed that the NHEJ-PD process generated by the two cuts occurs in different cell types, in primary and in immortalized cell lines, in different species and using different Cas9 proteins (either spCas9 or saCas9). Thus, the process seems to be mainly dependent on the NHEJ mechanism.

The combination of NHEJ-PD and the design of gRNAs to drive the activity of the Cas9 protein to very specific genome sites allow the prediction and generation of genotypes that could resemble very precisely mutations already described in humans. Here, we selected a specific region in exon 9 that generated mutations in the same location that a cluster of genetic mutations founds in PKD patients. This procedure is 10-times more efficient that the introduction of specific mutations by means of Homology Directed Repair and has allowed us to apply it to primary human hematopoietic progenitors. This procedure allows then to generate primary cells carrying almost similar human mutations in normal cells from healthy donors for those samples that are difficult or not ethically justified, and that could be used as cellular models for testing new available therapeutic strategies. Also, NHEJ-PD allow the selection of concrete sites which alterations could generate stop codons nearby, facilitating the generation of complete knock-out cellular models in a more efficient way. Moreover, we demonstrated that the same strategy applied to animal models is also very efficient.

Previous evidences have suggested that a minimum distance between the two cutting sites is required for the optimal NHEJ-PD. We have observed a similar requirement with an almost avoidance of precise deletion when cutting site guides were designed 8 bp apart. In addition, we demonstrate that also the orientation of the guides, and consequently the orientation of the Cas9 enzyme, is also important to allow optimal NHEJ-PD. A higher efficiency was obtained when both guides had the PAM sequence located in the sequence to be deleted, what we called PAM-in. The worst cutting efficiency was observed in the opposite orientation, PAM-out/PAM-out. Taking into account that the last 3 bp where common for both guides, steric interferences could be the responsible of this worse efficacy. Moreover, in those cases where PAM-in and PAM-out guides were used simultaneously, the PAM-in guide was more efficient than the PAM-out. Thus, factors like steric interactions but also orientation towards the sequence to be deleted are important for the efficiency of the CRISPR/Cas9 system to generate NHEJ-PD.

The use of guide pairs and the generation of NHEJ-PD has been also proposed as a therapeutic tool to precisely eliminate specific sequences that allowed the restoration of an almost completely functional protein. Mutations at dystrophin gene have been precisely eliminated by removing the sequence between exons 45-55 with the use of two guides restoring the function, and the Duchenne muscular dystrophy phenotype(Ousterout et al. 2015). The same strategy has been followed in epidermolysis bullosa, a recessive disorder caused by frameshift mutations at COL7A1 gene. The excision of the mutated sequences at exon 80 restores the collagen function and recovers a normal phenotype in keratinocytes(Bonafont et al. 2019). Our results clearly support this strategy and points out the importance of the design of the strategy in terms of CAs9 positioning.,

A potential drawback of the use of two guides is that the amount of off target mutations is multiplied by two. The optimization of the on line tools for the design of guide RNAs and the use of better systems to characterize them will help to improve their use.

In summary, we demonstrate here that the use of guide pairs in combination with the Cas9 nuclease generate NHEJ-PD in adult cells that facilitate the generation of cellular and animal models of specific mutations found human diseases. Moreover, these combinations could be used as a therapeutic tool for the treatment of genetic diseases where the excision of a defined sequence restores the function of the modified protein.

## ACKNOWLEDGEMENTS

The authors would like to thank Miguel A. Martin for the careful maintenance of NSG mice, and Mrs Aurora de la Cal, María del Carmen Sánchez, Soledad Moreno, Nadia Abu-sabha, Montserrat Aldea and Mr Sergio Losada for their dedicated administrative help. This work was supported by grants from “Ministerio de Ciencia, Innovación y Universidades y Fondo Europeo de Desarrollo Regional (FEDER)” (SAF2017-84248-P), “Fondo de Investigaciones Sanitarias, Instituto de Salud Carlos III” (Red TERCEL;RD16/0011/0011) and Comunidad de Madrid (AvanCell, B2017/BMD-3692). The authors also thank Fundación Botín for promoting translational research at the Hematopoietic Innovative Therapies Division of the CIEMAT. CIBERER is an initiative of the “Instituto de Salud Carlos III” and “Fondo Europeo de Desarrollo Regional (FEDER)”.

